# Automatic brain categorization of discrete auditory emotion expressions

**DOI:** 10.1101/2022.11.09.515555

**Authors:** Siddharth Talwar, Francesca M. Barbero, Roberta P. Calce, Olivier Collignon

## Abstract

Seamlessly extracting emotional information from voices is crucial for efficient interpersonal communication. However, it remains unclear how the brain categorizes vocal expressions of emotion beyond the processing of their acoustic features. In our study, we developed a new approach combining electroencephalographic recordings (EEG) in humans with an oddball frequency tagging paradigm to automatically tag neural responses to specific emotion expressions. Participants were presented with a periodic stream of heterogeneous non-verbal emotional vocalizations belonging to five emotion categories (Anger, Disgust, Fear, Happiness, Sadness) at 2.5 Hz. Importantly, unbeknown to the participant, a specific emotion category appeared at an oddball presentation rate at 0.83 Hz that would elicit an additional response in the EEG spectrum only if the brain discriminates the target emotion category from other emotion categories and generalizes across heterogeneous exemplars of the target emotion category. Stimuli were matched across emotion categories for harmonicity-to-noise ratio, spectral center of gravity, pitch, envelope, and early auditory peripheral processing via the simulated output of the cochlea. Additionally, participants were presented with a scrambled version of the stimuli with identical spectral content and periodicity but disrupted intelligibility. We observed that in addition to the responses at the general presentation frequency (2.5 Hz) in both intact and scrambled sequences, a peak in the EEG spectrum at the oddball emotion presentation rate (0.83 Hz) and its harmonics emerged in the intact sequence only. The absence of response at the oddball frequency in the scrambled sequence in conjunction to our stimuli matching procedure suggests that the categorical brain response elicited by a specific emotion is at least partially independent from low-level acoustic features of the sounds. Further, different topographies were observed when fearful or happy sounds were presented as an oddball that supports the idea of different representations of distinct discrete emotions in the brain. Our paradigm revealed the ability of the brain to automatically categorize non-verbal vocal emotion expressions objectively (behavior-free), rapidly (in few minutes of recording time) and robustly (high signal-to-noise ratio), making it a useful tool to study vocal emotion processing and auditory categorization in general in populations where brain recordings are more challenging.

## 1. Introduction

In humans and other animals, efficient categorization of emotion expressions is crucial for effective social interactions and survival. In Darwin’s milestone book “*The Expression of the Emotions in Man and Animals”* published in 1872, attention was mostly focused on the face as the carrier of emotion expressions. Since then, research on facial emotion expressions has led to the suggestion that at least six basic emotions-Anger, Fear, Disgust, Happiness, Surprise and Sadness (Ekman, 1993) can be expressed using specific facial movement (the FACS - Facial Acting Coding System Ekman & Friesen, 1978; Waller et al., 2020) and are thought to sustain different survival functions (Ekman, 1992; Panksepp, 1982). These emotion categories express similarly across different cultures (Elfenbein & Ambady, 2002), arise very early in development (Flom & Bahrick, 2007; Poncet et al., 2022), and can be found in our evolutionary ancestors (Darwin, 1872; Waller & Micheletta, 2013).

Although Darwin mentioned the importance of vocalizations as a carrier of affective signals (Darwin, 1872), it was suggested that the voice has yet to be proven a source for discrete emotion categories like the face (Ekman, 2009). Mounting evidence shows however that like the face, discrete emotion expressions can be delivered and decoded through vocal expressions with a high accuracy in humans (Cornew et al., 2010; Falagiarda & Collignon, 2019; Pell & Kotz, 2011). In fact, the same discrete emotion categories as those found for faces (Ekman, 1993) also express similarly through voices across cultures (Juslin & Laukka, 2003; Sauter et al., 2010; Sauter & Eimer, 2010; Scherer et al., 2001) and develop early in infancy (Izard et al., 1980; Mehler et al., 1978; Zhao et al., 2021).

Functional Magnetic Resonance Imaging (fMRI) studies have shown that common brain regions are activated across a wide range of emotion categories, for instance the amygdala (Fecteau et al., 2007; Wiethoff et al., 2009), the medial prefrontal cortex (Etkin et al., 2011; Kober et al., 2008) and ‘emotion voice areas’ in the auditory cortex (Ethofer et al., 2012). Despite the existence of these regions involved in the processing of various emotion expressions, it was also suggested that discrete emotions may recruit separate brain portions (Calder et al., 2001; Ethofer et al., 2009; Frühholz & Grandjean, 2013a; Johnstone et al., 2006; Kober et al., 2008; Kotz et al., 2013; Mauchand & Zhang, 2022; Phan et al., 2002; Phillips et al., 1998; Vytal & Hamann, 2010). However, it is also known that the distinct acoustic features that characterize different discrete categories of vocal emotions activate different patches in the core region of the auditory cortex that is known to be tonotopically organized (Talavage et al., 2004). Indeed, imaging studies revealed that the primary and secondary auditory cortices that typically process acoustic parameters of sounds, also show distinct activations for diverse affective vocalizations (Frühholz & Grandjean, 2013b). Acknowledging the fact that it was claimed that categorical response to vocal expression of emotions can be independent from acoustic amplitude and frequency cues (Giordano et al., 2021; Grandjean et al., 2005), whether the responses to emotional utterances and categorization processes are driven by low-level acoustic properties is still unclear.

Affective vocalizations have also been studied extensively using Event Related Potentials (ERPs). Responses as early as ∼ 100 ms were found to be modulated by emotional utterances compared to neutral vocalizations (Jessen et al., 2012), which may suggest that the differences are driven by variability of acoustic features (Salvia et al., 2014). Additionally, there are also evidences of enhancement of later ERP components such as the early posterior negativity (EPN: 200-350 msec) and late positive potential (LPP: ∼400 msec) from emotional utterances contrasted with neutral vocalizations (Frühholz et al., 2011; Jessen & Kotz, 2011). These later differences are thought to index the mechanism of affective categorization (Schirmer & Kotz, 2006) that may arise or be independent of acoustic differences. However, in most ERP studies, emotion related responses are compared to neutral, thus leaving unanswered whether the observed differences are due to static and dynamic acoustic cues such as fundamental frequency, intensity and temporal-spectral profile (Banse & Scherer, 1996; but see Bostanov & Kotchoubey, 2004; Pell et al., 2015).

In this study, we aimed to develop a novel approach combining electroencephalographic recordings (EEG) in humans with an oddball frequency tagging paradigm to provide an objective measure of automatic categorization of vocal emotion expressions beyond the processing of the stimuli acoustics. The frequency tagging approach (Regan, 1989) relies on the fact that under external stimulation of periodic stimuli, the brain areas that encode the stimuli synchronize at the exact same frequency (Norcia et al., 2015). We adapted the Fast Periodic Auditory Stimulation (FPAS, Barbero et al., 2021) to present different exemplar sounds of discrete emotion categories periodically such that the brain will elicit a response at the general rate of sound presentation (e.g. 2.5 Hz and its harmonics) if it can segregate different affective sounds. Importantly, within the sequence of affective vocalizations, there is also embedded another periodically occurring oddball emotion category (e.g. different exemplars of Fear are presented at every third position at 0.83 Hz). The brain will only respond at the oddball frequency and itss harmonics if it can *discriminate* the vocalization of the oddball target category from sounds of other frequent non-target categories as well as *generalize* all oddball vocalizations to one common emotion category (see Barbero et al., 2021).

From a fundamental point of view, our study aimed to investigate whether the human brain categorizes discrete emotions partially independently of the acoustic properties of the affective vocalizations. This was implemented by careful selection of non-verbal vocalizations with similar acoustic properties (spectral center of gravity, harmonicity to noise ratio, pitch) and introducing a second control stimulation sequence of frequency scrambled sounds with similar spectra-temporal profile (frequency content, sound’s envelope) and identical periodic constraints but disrupted intelligibility (Barbero et al., 2021; Dormal et al., 2018). We also conducted behavioral experiments to confirm the authenticity of the chosen emotion sounds and the unintelligibility of the scrambled stimuli. In addition to the frequency domain analysis, we conducted time lock analyses to investigate the time course of the response underlying emotion categorization. Finally, our study also aimed to provide a powerful tool to investigate the brain’s ability to categorize auditory emotion expressions objectively (the response lies at predefined frequencies of interest), robustly (with a high signal-to-noise ratio), and does not need any overt attention from the participant (behavior-free).

## 2. Materials and Methods

### 2.1 EEG Experiment

#### 2.1.1 Participants

Twenty-four participants (12 females, mean age: 22.29, S.D.: 2.33, range: 19-28 years) participated in the study. All participants had no history of neurological or audiological disorders and were right-handed. The experiment was approved by the local ethical committee of the Catholic University of Louvain (UCLouvain, project 2016-25). All participants provided written informed consent and received financial compensation for their participation.

#### 2.1.2 Stimuli

We selected sounds of five primary emotion categories: Anger, Disgust, Fear, Happiness and Sadness. The stimuli were extracted from clips in which professional actors and actresses performed emotion expressions in varied styles and intensities without any linguistic content (Belin et al., 2008). Additionally, a few stimuli were selected from a database of non-verbal vocalizations depicting several distinct varieties of emotion categories (Cowen & Keltner, 2017). 96 heterogenous sounds were cropped to a length of 350 ms to allow periodic presentation. Root mean squared values of all sounds were equalized and 10 msec ramps were applied at the start and end of the stimuli to avoid clicking. A potential confound in this study of processing of vocal emotion responses is that the acoustic properties of sounds could partially account for the categorical response to a specific emotion, if not accounted for. We therefore confirmed that the sounds have comparable spectral center of gravity (COG; F = 1.1882, p = 0.3224, ηp2 = 0.0567), harmonicity-to-noise ratio (HNR; F = 1.6930, p = 0.1599, ηp2 = 0.0789) and pitch (F = 1.7618, p = 0.1448, ηp2 = 0.0819) across the five emotion categories (Figure 1a). The acoustic properties were computed using custom scripts in Praat (Boersma & Weenink, 2001).

**Fig 1.**
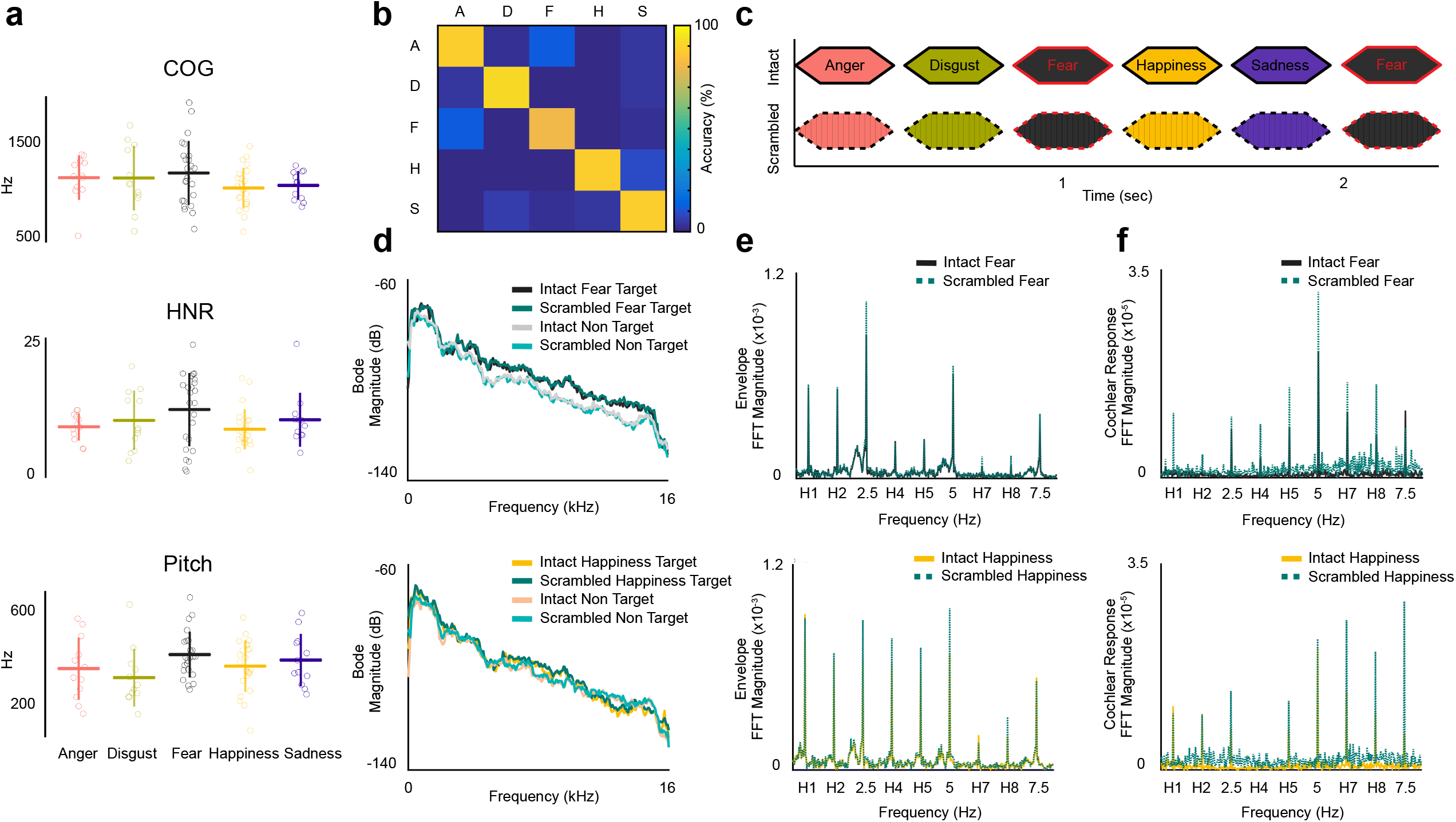
Stimuli and sequences: A) The selected sounds of five emotion categories have similar spectral properties: center of gravity (COG), harmonicity to noise ratio (HNR) and pitch. B) Behavioral experiment validated 84 short non-verbal vocalizations to depict the appropriate emotion category. A=Anger, D=Disgust, F=Fear, H=Happiness, S=Sadness. C) Sounds were presented periodically at 2.5 Hz. The target emotion (e.g. fear in this illustration) repeated at every third position leading to a target presentation rate of 0.83 Hz while other emotion categories were presented randomly. D) Bode plot shows similar averaged spectral power of intact and scrambled stimuli for target and non-target sounds, for each emotion condition. E) Similar averaged FFT distribution of the envelope between intact and scrambled sequences indicates similar temporal profiles across the two types of sequences in each emotion condition. F) No significant difference found in the averaged FFT magnitudes of the simulated cochlear response to intact and scrambled sequence, verifying similar spectro-temporal profile of all sequences as well as the processing of acoustic features at the cochlear level

Subsequently, we verified whether the resultant sounds could be categorized accurately across participants (same emotion category as the actors tried to emote). We conducted a behavioral experiment where 10 participants (who did not participate in the EEG experiment) categorized each sound to an emotion category in a five-alternative forced choice task. 3 blocks each composed of 96 stimuli with 1 repetition per block were presented in a randomized fashion. Finally, 84 sounds across 5 categories were selected (Anger, Disgust, Sadness: 12 each; Fear, Happiness: 24 each) indicating 80% or more accuracy across all participants for an emotion category, which were then used to build sequences (Figure 1b). We required double the number of stimuli for Fear and Happiness categories to present an equal number of unique stimuli of each category in the sequence (More details in section 2.1.3). Each emotion category consisted of an equal number of sounds by male and female actors, to incorporate heterogeneous spectra-temporal profiles across the stimuli set.

Furthermore, the selected stimuli were frequency scrambled using the method described in Dormal et al., 2018. This method preserves the overall frequency content of the original sounds while disrupting their harmonicity and intelligibility. Specifically, we applied a Fast Fourier Transform (FFT) to each sound and shuffled the magnitude and phase of frequency bins within consecutive windows of 200 Hz. After implementing the inverse FFT to compute the scrambled sounds, we applied the original envelope to the resultant scrambled waveforms by extracting the envelope from the respective original sounds using the Hilbert transform. This procedure led to a disruption in harmonicity and intelligibility of the scrambled sounds (as confirmed in a behavior experiment: see section 3.3), while keeping their spectral-temporal structure almost identical to the original / intact sounds (Figure 1d, 1e). The overall energy (root mean square: RMS) of all scrambled sounds were equalized and ramps of 10 msec were applied, same as frequency intact sounds.

#### 2.1.3 Sequences and Procedure

The affective sounds were placed together to prepare the periodic sequences with an inter stimulus interval (ISI) of 50 msec to facilitate sound segregation and discrimination. Thus, the general rate of presentation or base rate was set at 2.5 Hz (1000/(350+50) msec). Importantly, each third sound in the sequence belonged to a specific emotion category, which was presented periodically at a target frequency of 0.833 Hz (2.5 Hz / 3 sounds), while all sounds of non-target categories were presented non-periodically. Two emotion conditions were created, where Fear and Happiness were used as the target category, respectively. Fear was chosen due to its salience and evolutionary function (Stanley, 1984). Happiness was selected due to its differentiated valence amongst other chosen ‘negative’ valence emotions. For each emotion condition, different sequences were constructed such that the order of stimuli in each sequence was randomized to increase generalization. The order of each frequency intact sequence was replicated in a frequency scrambled sequence. As a result, each intact sequence had an identically ordered control scrambled sequence (Figure 1c). With 24 targets of 1 emotion category (Fear or Happy) and 48 non-targets across other 4 emotion categories in any sequence, each sound was presented twice in a sequence (total sounds: 144) leading to a length of 57.6 seconds. The sequence included 2 seconds of fade-in and fade-out, where the volume gradually increased from 0 to max and decreased from max to 0 respectively, to avoid abrupt movements by participants that cause artifacts in the EEG data.

For EEG acquisition, participants sat in a dimly lit room and were presented two different sequence conditions (intact sequence with conditions-Fear and Happiness as oddball) and to control for low-level acoustic confounds (scrambled frequency sequences), leading to four groups of sequences: Fear Intact, Fear Scrambled, Happiness Intact, Happiness Scrambled. Ten instances of each sequence group (total: 40) were created and presented to each participant in a pseudo-randomized fashion. The participants kept their eyes closed during the length of the sequence and were instructed to press a button when they heard a sound lower in volume compared to other sounds. The lower volume was implemented by reducing the sounds’ root mean square (RMS) value by a factor of 10. This attention target was presented 6 times per sequence in a randomized fashion, excluding the 2 seconds fade-in and fade-out at the start and end of the sequence. The experiment was designed on MATLAB R2016b (Mathworks) with Psychtoolbox and extensions (Brainard, 1997; Kleiner et al., 2007; Pelli, 1997).

#### 2.1.5 EEG Acquisition

The speakers were placed at a 1 m distance from the participants. EEG was acquired with a Biosemi ActiveTwo System (https://www.biosemi.com/products.htm) using 128 channel Ag/AgCl electrodes. International 10-20 system were used for the recording sites as well as their intermediate positions (position coordinates can be found at https://www.biosemi.com/headcap.htm). Additionally, two surface electrodes were applied at mastoids. The acquisition sampling rate was 512 Hz. Impedances of all electrodes were kept below ±25 mV.

#### 2.1.6 Analysis

Data was analyzed using Letswave 6 (https://github.com/NOCIONS/Letswave6) and Fieldtrip toolbox (Oostenveld et al., 2011) on MATLAB R2016b (Mathworks), and using custom built scripts in MATLAB and Rstudio (RstudioTeam, 2015).

##### 2.1.6.1 Preprocessing

The pipeline for preprocessing and frequency domain analysis has been replicated in many frequency tagging studies in audition (Barbero et al., 2021) and vision (Bottari et al., 2020; Dzhelyova et al., 2017; Retter & Rossion, 2016; Rossion et al., 2015, 2020; Volfart et al., 2021). A fourth order Butterworth bandpass filter was applied on raw continuous EEG data from 0.1 to 100 Hz, along with a notch filter at 50 and 100 Hz of a width of 0.5 Hz to attenuate the power line noise. The data was then downsampled to 256 Hz to facilitate data handling and storage. Next, data was segmented from 2 seconds before the onset of the sequence and 2 seconds after the end of the sequence resulting in a trial length of 61.6 seconds. Subsequently, the ICA matrix of each segmented trial was computed using RUNICA (Makeig et al., 1995) and the components were visually inspected. Only the noisy frontal components due to facial movements were taken into consideration for deletion, and at most one component was deleted in 18 out of a total 24 participants. Visual inspection of resultant trials was conducted, and an average of 1.6 noisy trials were deleted out of the total 40 trials per subject (at most, 5 trials were deleted for 2 subjects). Additionally, we implemented linear interpolation using 3 closest neighbors for noisy channels. We interpolated 2 channels for 5 subjects (FT8, FC5; T8h, AF7; FPz, AFF2; AF3, P9; PPO5, AFF2) and 1 channel for 4 other subjects (I1; I1; C6; CPP5h). The trials were subsequently re-referenced to average reference and divided into two emotion conditions (Fear and Happiness) and its two sequence types (Intact and Scrambled)-Fear Intact, Fear Scrambled, Happiness Intact and Happiness Scrambled.

##### 2.1.6.2 Frequency Domain Analysis

Considering the presentation rate of the target emotion category and the frequency resolution (1/duration of the sequence), all trials were re-segmented from 2 seconds after onset (to remove fade-in) to a length of 52.8 seconds, to contain an integer of target presentation cycles (0.833 Hz). Following, we grand averaged across all participants, all the trials per each emotion condition and sequence type to attenuate the noise and signals irrelevant to the experiment (brain processes not relevant to stimuli presented). Consequently, a Fast Fourier Transform (FFT) was applied on the averaged trials. Amplitude spectra extended from 0 to 128 Hz with a frequency resolution of 0.0189 Hz, allowing us to isolate the base response (general presentation rate) at 2.5 Hz and target response at 0.833 Hz along with their respective harmonics. The harmonic responses in addition to fundamental frequency (in this case 2.5 Hz or 0.83 Hz) can be accounted for in relation to complex responses of the brain, corresponding to the principles of frequency domain analysis of periodic signals (Retter et al., 2021). Thus, we applied a criteria to choose the harmonics that were required in further analysis since the summation of harmonics in the FFT can indicate the overall responses in single values (Retter et al., 2021; Retter & Rossion, 2016). Significance of harmonics was determined by first pooling all channels and computing the z-score at each harmonic bin, considering 12 bins at each side of the frequency bin of interest, excluding the immediate adjacent bins and the maximum and minimum of the entire window (Retter & Rossion, 2016). For each target emotion condition (Fear and Happiness) and sequence type (intact and scrambled), consecutive harmonics that displayed a z-score > 2.32 (p<0.01, 1-tail, signal>noise) were considered as significant. We excluded the base frequency bins (2.5, 5, 7.5 Hz etc) while observing consecutive frequency bins for target harmonics. The chosen number of consecutive harmonics of intact and scrambled sequence types were equalized by considering the highest number of significant consecutive harmonics in any of the two types of sequences, acknowledging that adding responses at non-significant harmonics is not detrimental (i.e. adding zeros, Retter et al., 2018). Further, to quantify the responses at base and target frequencies independently, we computed the baseline subtracted magnitude from the FFT of each condition and type, where the window for this calculation was defined similarly to the computation of z-scores (12 bins on either side except maxima, minima and adjacent bins). The baseline subtraction takes into account the different noise profiles across different frequency bands (Luck, 2014), especially higher noise at low frequencies (<1Hz) in EEG recordings. Consequently, we summed the chosen significant harmonics to finally represent them as topographical maps.

To find the electrode of interest involved in emotion categorization for Fear and Happiness that also could be independent of low-level processes, we subtracted the grand averaged FFT spectra of frequency intact sequence and frequency scrambled sequences (*intact - scrambled*) for each target emotion condition separately. Then we extracted chunks of the resultant spectra, where each chunk consisted of 25 frequency bins and the 13^th^ bin included the chosen harmonic (12 neighboring bins on either side of the harmonic to define noise). The number of chunks was defined as the number of chosen harmonics in each emotion condition. The mean of the chunks was computed (sum/number of chunks) and the z-score was computed at the harmonic (13^th^ bin) for each channel (Volfart et al., 2021). Finally, we performed FDR (Benjamini & Hochberg, 1995) correction across 128 channels for multiple comparisons, for each emotion condition.

##### 2.1.6.3 Time Domain Analysis

The raw data was filtered through the Butterworth bandpass and the notch filter with the same parameters as in frequency domain analysis. Additionally, another notch filter was applied at the base presentation frequency and harmonics (2.5, 5, 7.5 Hz), to remove responses elicited at the base rate by general auditory processes not linked to target emotion discrimination. Stimulus-locked epochs of 800 ms ranging from the onset of one stimulus prior to the target stimulus to the end of the target stimulus were extracted such that each sequence yielded 44 epochs. The first 2 and last 2 epochs were deleted due to fade in and fade out of the sequence. Noisy epochs with amplitude exceeding ±100µV in any channel were deleted. The resultant epochs were re-referenced to averaged mastoids, then equalized across conditions and types separately and averaged for each subject. Baseline correction was implemented by subtracting the signal within 400 ms pre-stimulus activity, corresponding to 1 cycle of base rate, i.e. the prior epoch to the target epochs (Dzhelyova et al., 2017). Finally, the baseline subtracted epochs were grand averaged across all subjects for each condition and type separately.

To compare the temporal activity of emotion categorization, we conducted a timepoint-by-timepoint, one-tailed t-test between frequency intact and scrambled sequence types across subjects for each electrode in each emotion condition. Segments of data were considered significantly different only if the two conditions (intact vs scrambled) were different for more than 25 ms, i.e. >13 consecutive time-points (Chen et al., 2021). For each emotion condition, FDR correction was applied across 128 channels to correct for multiple comparisons.

### 2.2 Cochlear Model and Envelope

In this study, we aimed to find an EEG response, if any, specific to an emotion category at 0.833 Hz and its harmonics (1.66, 3.33 etc), which may at least be partially independent from responses caused due to low-level acoustic features, by contrasting frequency intact and scrambled sequences. Thus, to validate whether the differences in temporal structure of the target emotion category and non-target emotion categories don’t account for the EEG response, as in a case of an oddball paradigm, we compared the temporal envelopes of intact and scrambled sequences. First, we extracted the envelope of each sequence using the Hilbert Transform. Then, we computed the FFT of all envelopes and averaged them across each sequence type (intact and scrambled) for each participant. Further, we chose the harmonics of interest in the FFT based on the empirical EEG data (see section 2.1.6.2 for the procedure and section 3.1.2 for EEG results) and summed the FFT magnitudes across the chosen bins of interest for every subject. Finally, a 1-tail t-test was calculated across all subjects to contrast intact from scrambled sequences. This procedure was performed for Fear and Happiness emotion conditions separately.

Furthermore, we assessed how spectral and temporal characteristics of sound alone can influence early responses processed in the cochlea before moving up the auditory pathway towards the sensory cortices. For this reason, we employed gammatone filters using Auditory Toolbox (Slaney, 1994) to simulate a cochlear response to a given sequence. The FFT was applied to the simulated cochlear responses that were computed from all sequences prepared for each EEG participant, and were further averaged for each sequence type and condition. We chose the harmonics of interest in the FFT of the cochlear response based on the empirical EEG data (see section 2.1.6.2 for the procedure and section 3.1.2 for EEG results) and summed the FFT magnitudes across the identified bins of interest for each subject. Lastly, we compared the summed FFT magnitudes of intact and scrambled sequences across all subjects using a 1-tail t-test (intact > scrambled) for each condition separately.

### 2.3 Behavioral Experiment

We conducted a behavioral experiment to evaluate whether the participants could efficiently categorize emotion from a stream of short bursts of affective voices and classify them to discrete emotion categories, and to confirm the unintelligibility of the scrambled sounds.

#### 2.3.1 Participants

21 out of 24 participants (11 women, age: 22.28, SD: 2.36, range: 19-28 years) who took part in the EEG experiment participated in the behavioral experiment. This session was scheduled after the EEG experiment to avoid familiarity of the sounds during the EEG session. All participants had no history of neurological or audiological problems and were all right-handed. The experiment was approved by the local ethical committee of Catholic University of Louvain (project 2016-25). All participants provided written informed consent and received financial compensation for their participation.

#### 2.3.2 Procedure

Stimuli from the EEG session were used in the behavioral experiment (see section 2.1.2). The behavioral session was divided into two tasks: 1) a “Sequence task” where subjects were asked to identify a target emotion category amongst sounds presented in a short sequence and 2) an “Isolation task” where each sound was presented separately. The sequence task was divided into four blocks where each block consisted of 48 short sequences, each of five emotional sounds. Two of the four blocks required the participants to identify if a Fearful vocalization was present amongst all the sounds presented, while the other two blocks asked them to identify if a Happy expression was present in the sequence. To mimic the structure of sequences used in EEG, we inserted the target emotion at the third place in each short sequence. Subjects were asked to perform a two-alternative forced-choice task (target emotion present or not) with half of the sequences in each block (24/48) consisting of the target emotion category (Fear or Happiness according to the block). All sequences and blocks were presented randomly. Later, participants performed the “Isolation task” which was also divided into four blocks each consisting of 84 sounds. In two blocks, subjects were asked to listen to a single affective vocalization and answer three questions- a) classify the sound to one of the five emotion categories – Anger, Disgust, Fear, Happiness or Sadness in a five-alternatives forced choice task. b) rate the valence of each sound on a scale from 1 to 5 (1 = most negative, 3 = neutral, 5 = most positive). c) rate the arousal evoked by the stimuli on a scale from 1 to 5 (1 = not aroused at all, 5 = most aroused). In the other two blocks, the scrambled version of the same sounds were presented and the subjects were asked to classify them to an emotion category in a five-alternatives forced choice task. All sounds and blocks were presented in a random order. The experiment was implemented on MATLAB R2016b (Mathworks) with Psychtoolbox and extensions (Brainard, 1997; Kleiner et al., 2007; Pelli, 1997).

## 3. Results

### 3.1 Experiment 1- EEG

We anticipated similar responses at the general rate of presentation / base frequency and its harmonics (i.e., at 2.5, 5, 7.5 Hz etc) when both frequency intact and scrambled sequences were presented, indicating segregation of all sounds at the presentation rate as well as low-level acoustic processing. Importantly, we predicted to observe an enhanced response at the target frequency and its harmonics (i.e. at 0.833, 1.666, 3.333, 4.166 Hz) for the frequency intact but not for scrambled sequence, signifying emotion categorization beyond acoustic properties. We built two conditions of sequences: Fear and Happiness, to investigate differences, if any, in the EEG responses to these discrete emotion categories. Additionally, we conducted time domain analysis to investigate the time course of the response underlying emotion categorization.

#### 3.1.1 Frequency Domain: Base Response

2 consecutive significant harmonics for Fear (2.5, 5 Hz) and 3 for Happiness were identified (2.5, 5, 7.5 Hz). We observed a similar base response at the general rate of presentation of sounds and its harmonics for both Fear and Happiness emotion conditions, and between intact and scrambled types. The similar scalp topographies of the sum of baseline subtracted magnitudes calculated at these frequencies suggest that different vocalized emotional sounds and the frequency scrambled unintelligible sounds were processed in a similar way across all sequences. (Figure 2a).

**Fig 2.**
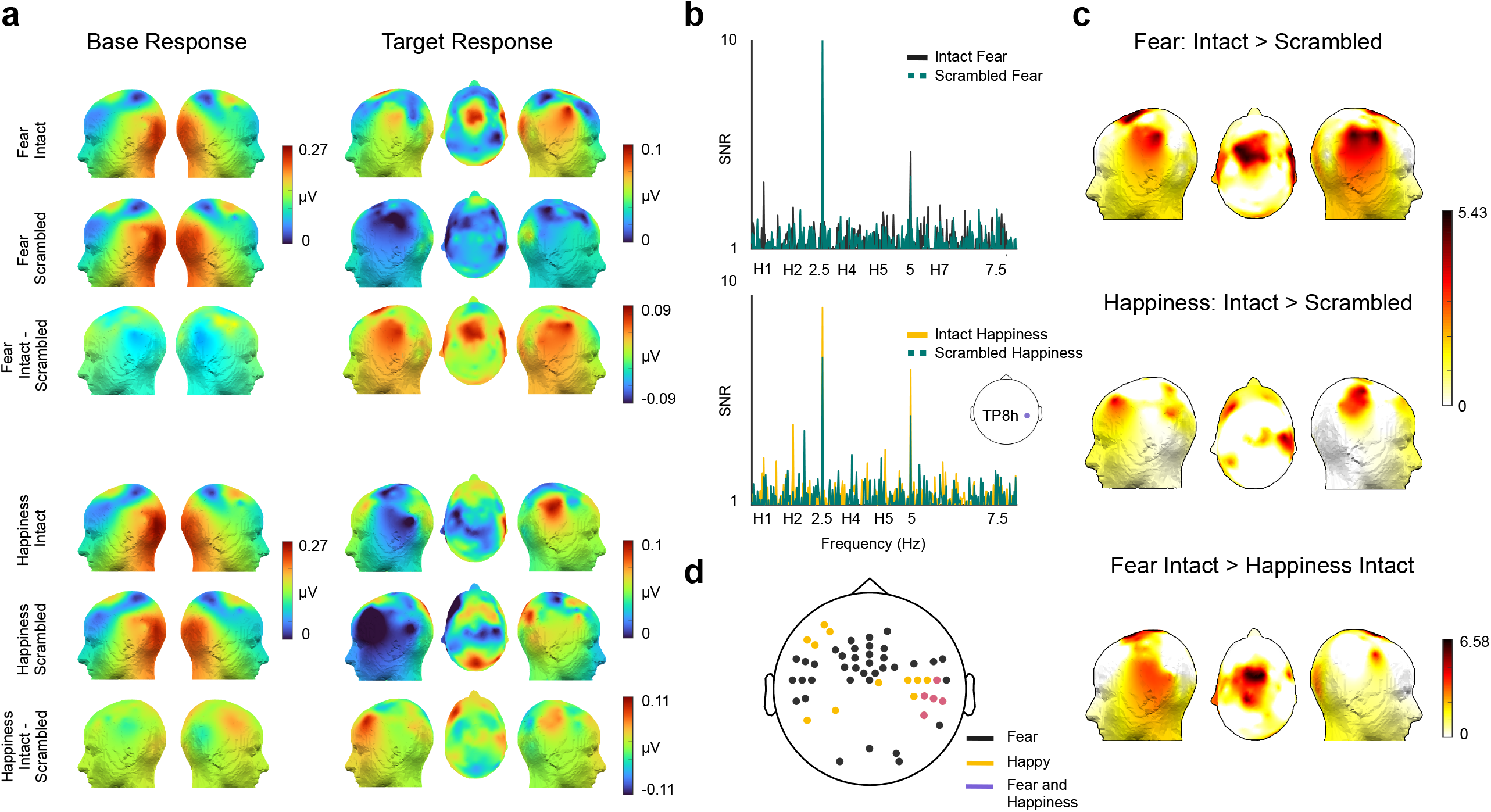
EEG results: A) Sum of baseline subtracted magnitudes are represented as topographies. The base response appears similar across all conditions and types. The target response to Intact Fear is localized to bilateral temporal and central areas while response to Intact Happiness is elicited in the right temporal and left frontal region. Higher responses observed for intact stimuli sequences than scrambled stimuli sequences. B) signal to noise ratio (SNR) at channel TP8h for both emotion conditions to visualize the response across frequencies. H1 and H*n* refer to the first harmonic of target frequency i.e. 0.83 Hz and higher harmonics, respectively. C) Scalp regions eliciting significant emotion selective responses. D) Distinct, yet overlapping channels selective to Fear and Happy

#### 3.1.2 Frequency Domain: Target Response

The target response was defined as the summation of consecutive significant harmonics, i.e. 5 harmonics for Fear (0.833, 1.666, 3.333, 4.166 and 5.83 Hz) and 4 for Happiness condition (0.833, 1.666, 3.333 and 4.166 Hz). The sum of baseline subtracted magnitudes of the harmonic bins of interest revealed a higher magnitude for intact sequence type in comparison to scrambled for both Fear and Happiness conditions (Figure 2a). We also computed the signal to noise ratio (SNR) using the same criteria to estimate the noise (12 bins on either side, excluding the immediate bins on either side, the maximum and minimum), at the right temporal channel TP8h for both conditions for visualization purposes (Figure 2b), showing higher SNR at the harmonics of interest for intact in comparison to scrambled.

To find the channels contributing to the responses at the target frequency and harmonics, we first computed the difference between the grand averaged Fast Fourier spectra of frequency intact and scrambled conditions at each electrode and computed the z-score for the 2 conditions: Fear and Happiness (see methods section 2.1.6.2). The one-sided p-values from the z-test (intact – scrambled > 0) were corrected for multiple comparisons using False Discovery Rate correction (FDR, Benjamini & Hochberg, 1995). We found 42 significant channels for Fear selective response in bilateral temporal areas (Figure 2c- maximum in right temporal: FT8h, z = 5.3219, p = 5.1345 × 10^−8^; in left temporal: TP7h, z = 4.7200, p = 1.1792 × 10^−6^) and fronto-central electrodes (maximum at FFC3, z = 5.4294, p = 2.8272 × 10^−8^). 16 significant channels for Happiness condition were clustered in right temporal (maximum at C6h, z = 4.8022, p = 7.8466 × 10^−7^) and left frontal area (maximum at AFF5, z = 4.3399, p = 7.1274 × 10^−6^). All other significant channels with their z-scores are tabulated in Table S1 in supplementary material.

We found five significant overlapping channels across Fear and Happiness conditions at the right temporal area, that may indicate elicitation from regions common to emotion processing, regardless of the category (P6, TP8h, TP8, CP6 and T8h, Figure 2d). However, we also observed different clusters of significant electrodes between both conditions. To investigate further whether some channels were more selective to Fear than Happiness or vice versa, we subtracted the FFT of Happiness intact from Fear intact, and computed the z-score at the bins of interest (See section 2.1.6.2). Since previously we selected different number of harmonics to quantify Fear (5) and Happiness (4) responses, we equalized the number of harmonics to the maximum i.e. 5 harmonics since the activity across non-significant harmonics to compute overall response is not detrimental (i.e. as adding zeros). A 2-tail t-test revealed 38 significant channels selective to Fear compared to Happiness (Figure 2e) mostly clustered at and around the central region (maximum at FFC1, z = 6.4061, p = 7.4645 × 10^−11^). We did not find any channels significantly more selective to Happiness than Fear.

#### 3.1.3 Time domain

We compared the grand averaged trials of intact and scrambled with a timepoint by timepoint 1-tail t-test separately for each emotion condition. The analysis revealed two significant windows for the Fear condition (Figure 3)-between 135 ms to 176 ms (CP6 at 152 ms: SD = 0.7978, t_(23)_ = 4.1631, p = 0.001 FDR corr, Cohen’s d = 3.8140) and between 311 ms to 398 ms (Cz at 339ms: SD = 0.8983, t_(23)_ = 4.5269, p = 8.8496 × 10^−4^ FDR corr, Cohen’s d = 4.1993). These differences expressed across central, frontal and parietal electrodes. Happiness intact trials elicited a stronger response than scrambled in only one late window: 319 ms to 356 ms (Cp6 at 339ms: SD = 0.5628, t_(23)_ = 5.0178, p = 0.0013 FDR corrected, Cohen’s d = 4.1993) expressing across right temporal channels. Thus, in addition to different topographic representations, we also found different temporal orders of categorization of Fear and Happiness, where EEG indexing processing of Fear occurred earlier than Happiness. Similar to frequency domain analysis, four channels in the right temporal region: P6, TP8h, CP6 and T8h were found to elicit responses significantly greater for intact sequences than scrambled for both Fear and Happiness conditions.

**Fig 3.**
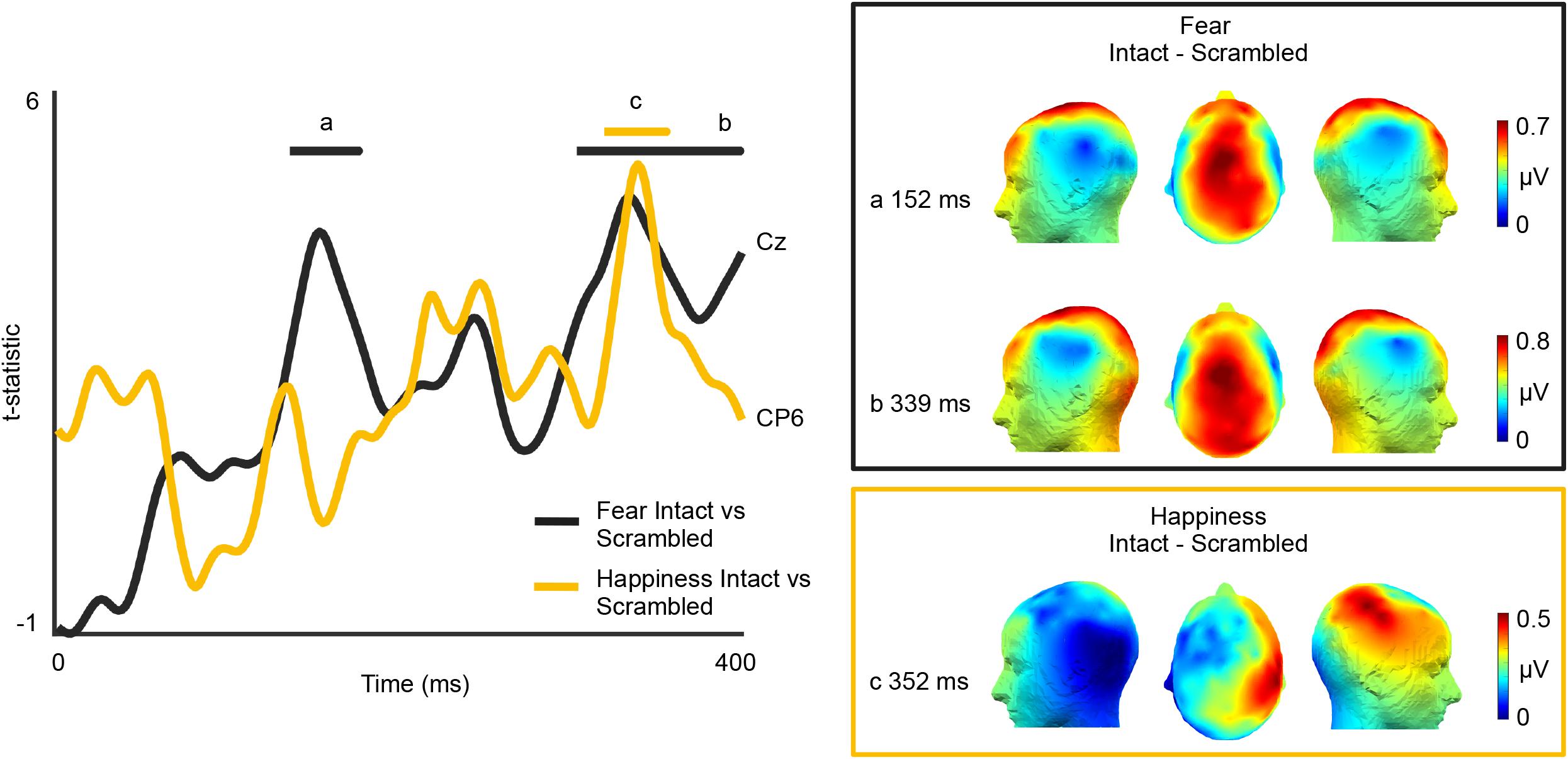
EEG time domain results: t-statistic of Fear intact > scrambled and Happiness intact > scrambled plotted with respect to time. The representation of Fear on contrasting with scrambled sequences, evolves early (peaking at 152 ms) and later (339 ms) across fronto-central-parietal channels; only one significant window for happiness was found peaking at 352 ms in the right temporal parietal region. The topographies represent the EEG amplitude at the peak time points

### 3.2 Cochlear Model and Envelope

After preparing the sequences of affective sounds for each subject, the FFT spectrum of the envelope of sequences of both emotion conditions (Fear and Happiness) and types (Frequency Intact and Scrambled) were averaged separately. Then, the FFT magnitude was summed at the harmonics (defined as per the empirical EEG results: 5 for Fear, 4 for Happiness) and contrasted with a 1-tail t-test (intact *>* scrambled) revealing similar temporal structure for both type of sequences (*Fear*: t_(23)_=-0.3718, p=0.6432, Cohen’s d=-0.1759; *Happiness*: t_(23)_=-0.7062, p=0.75644, Cohen’s d=-0.1442) (Figure 1e). Similarly, we computed a 1-tail t-test between the summed FFT magnitudes of harmonics of interest of the simulated cochlear response to the two types of sequences (intact *>* scrambled, *Fear*: t_(23)_=-6.9465, p=0.9999, Cohen’s d=-1.4180; *Happiness*: t_(23)_=-11.7324, p=0.9999, Cohen’s d=-2.3949), indicating similar processing of the acoustic cues at the cochlear level for both type of sequences, for each emotion condition (Figure 1f).

### 3.3 Experiment 2- Behavior

We conducted a separate behavioral experiment with the same participants who took part in the EEG experiment to validate: a) if participants could categorize a short non-verbal affective vocalization played in a sequence (Sequence task), b) the unintelligibility of the scrambled sounds (Isolation task). Sensitivity indices were calculated for each task using D-primes (d’) constituting an unbiased quantification of performance in detection tasks considering both hits and false alarms (Hautus et al., 2021; Tanner Jr. & Swets, 1954). In the Sequence task, all participants identified the presence of a target emotion amongst a stream of other emotional sounds well above chance, for both Fear, (one tail t-test d’>0 - d’ values mean = 1.3866, SD = 0.5116, t_(20)_ = 12.4212, p = 7.37 × 10^−11^, Cohen’s d = 3.83) and Happiness (one tail t-test d’>0 - d’ values mean = 2.2399, SD = 0.8638, t_(20)_ = 11.882, p = 1.61 × 10^−10^, Cohen’s d = 3.66; Figure 4a).

**Fig 4.**
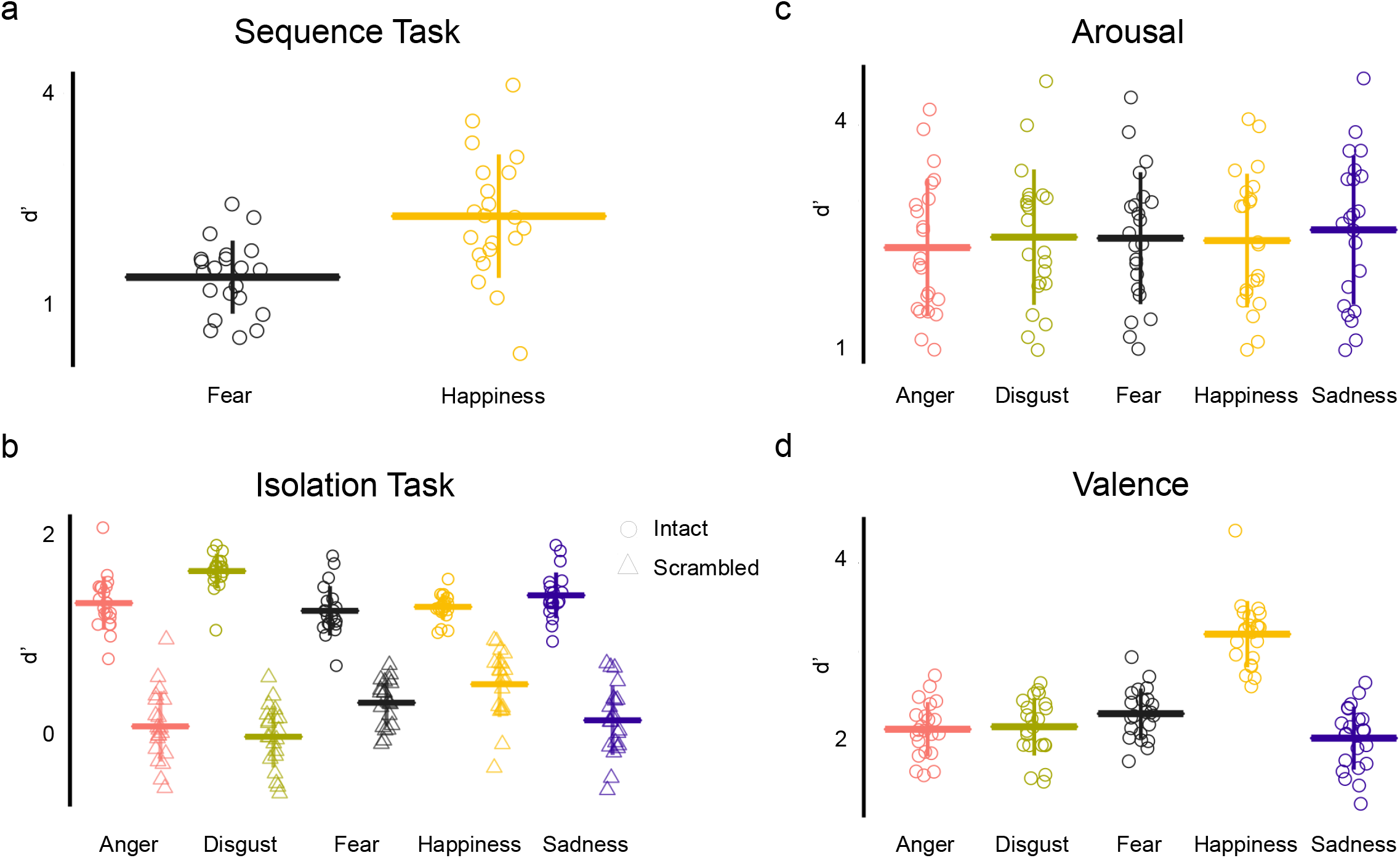
Behavioral results: A) Sequence task: Participants could identify the presence of an exemplar of the target emotion category amongst a short sequence of sounds well above chance level. B) Isolation task: Intact sounds were categorized to the appropriate emotion category, with worse performance for scrambled sounds, thus verifying the disruption in unintelligibility. C) Similar arousal ratings across all emotion categories suggest no involvement of arousal in the EEG responses. D) Happiness was the only ‘positive’ emotion amongst other categories. Greater values indicate positive valence

In the isolation task, all intact stimuli were correctly categorized above chance level (Figure 4b: one tail t-test d’>0 *Anger*: d’ values mean = 1.3214, SD = 0.2711, t_(20)_ = 22.3376, p = 1.33 × 10^−15^, Cohen’s d = 6.89; *Disgust*: d’ values mean = 1.6475, SD = 0.1768, t_(20)_ = 42.7111, p = 0, Cohen’s d = 13.1809; *Fear*: d’ values mean = 1.2428, SD = 0.2511, t_(20)_ = 22.6799, p = 9.99× 10^−16^, Cohen’s d =6.99; *Happiness*: d’ values mean = 1.2828, SD = 0.1309, t_(20)_ = 44.9205, p = 0, Cohen’s d = 13.86; *Sadness*: d’ values mean = 1.4006, SD = 0.2331, t_(20)_ = 44.9205, p = 0, Cohen’s d = 8.49). For scrambled stimuli, the categorization of three categories- Anger, Disgust and Sadness were at chance level (one tail t-test d’>0, *Anger*: d’ values mean = 0.0665, SD = 0.3587, t_(20)_ = 0.8501, p = 0.4053, Cohen’s d = 0.26; *Disgust*: d’ values mean = -0.0387, SD = 0.3132, t_(20)_ = -0.5670, p = 0.577, Cohen’s d = -0.1749; *Sadness*: d’ values mean = 0.1284, SD = 0.3523, t_(20)_ = 1.6704, p = 0.1104, Cohen’s d = 0.5155). However, d’ of scrambled versions of Fear and Happiness were found to be significantly higher than chance (one tail t-test d’>0, *Fear*: d’ values mean = 0.3071, SD = 0.2332, t_(20)_ = 6.0349, p = 6.71 × 10^−6^, Cohen’s d = 1.8624; *Happiness*: d’ values mean = 0.4956, SD = 0.3341, t_(20)_ = 6.7958, p = 1.31 × 10^−6^, Cohen’s d = 2.0972). Two explanations could be given for this result. First, there was a higher number of Fear and Happiness stimuli than other categories, since we chose the number of stimuli based on equal presentation of each sound in a sequence. Thus the higher d’ values could indicate an effect of training. Secondly, it is possible that the scrambled sounds of Fear and Happiness categories carried some amount of acoustic information which made them slightly recognizable. However, contrasting intact Fear vs scrambled Fear sounds, as well as intact Happiness vs scrambled Happiness stimuli indicated that the intact vocalizations carried significantly more information than their scrambled version (two tail t-test: *Fear:* SD = 0.2996, t_(20)_ = 14.306, p = 5.74 × 10^−12^, Cohen’s d = 3.12; *Happy*: SD = 0.3841, t_(20)_ = 9.3916, p = 8.99 × 10^−9^, Cohen’s d = 2.04)

We further analyzed the arousal and valence ratings of the intact sounds. We observed similar arousal levels across all the discrete emotion categories (ANOVA F = 0.15, p = 0.9612, ηp² = 0.5848; Figure 4c). We observed an effect of emotion on the valence ratings (Figure 4d: ANOVA F = 28.15, p = 6.8228 × 10^−16^, ηp² = 17.7659). As expected, post hoc t-tests revealed that valence ratings for Happiness were significantly different than other categories (Happiness vs Anger p = 2.98 × 10^−8^; Happiness vs Disgust p = 1.98 × 10^−8^; Happiness vs Fear p = 1.81 × 10^−8^; Happiness vs Sadness p = 5.96 × 10^−8^, FDR corrected).

## 4. Discussion

We demonstrate how combining electroencephalographic recordings (EEG) in humans with an oddball frequency tagging paradigm provides a robust and objective measure of a categorical brain response to short bursts of affective vocalizations. To disentangle the contributions of low-level auditory processing of affective vocalizations to representations of a higher order categorization process, we carried out a careful selection of sounds to match their spectra-temporal properties. We first matched the stimuli based on the main spectral features: spectral center of gravity, harmonicity-to-noise ratio (HNR) and pitch, such that no emotion category was spectrally and significantly different than another (Figure 1a). Next, sounds could be efficiently classified to a discrete category of emotion: Anger, Disgust, Fear, Happiness and Sadness, validated with a behavioral experiment (Figure 1b). Then, we introduced a sound sequence with identical periodic constraints like the intact sequences but scrambled sounds such that their spectra-temporal structure are similar but their intelligibility and harmonicity are disrupted (Barbero et al., 2021; Dormal et al., 2018, Figure 1c,d,e). To verify whether the envelopes of the sounds alone could elicit the EEG emotion response, we compared the FFT magnitude of the extracted envelopes of frequency intact and frequency scrambled sequences to find no significant difference at the frequency bins of interest (Figure 1e). As a final validation, we also used Gammatone filters to simulate the cochlear response to intact and scramble sequences and showed that there is no difference at the level of early peripheral auditory processing (Figure 1f). Thus, any difference in the EEG response to intact versus scrambled sequence cannot be simply explained by a cochlear simulation of acoustic response, nor by their spectral or envelope structure. Altogether, our stimuli selection procedure and control analyses assert that the observed categorical response to specific emotion expression categories is at least partially independent from low-level acoustic features and therefore reflects a higher-level categorization process.

We relied on a frequency tagging technique which allowed us to isolate the responses objectively and automatically at specific known frequencies without having the participants to overtly respond to the sounds, thus avoiding the decisional and attentional processes to contaminate the EEG response (Levy et al., 2003). Although the frequency of presentation of sounds of the target emotion category (0.83 Hz) lies at the lower spectrum of EEG which is susceptible to noise (Luck, 2014), we acknowledged the trade-off between the frequency of presentation of target emotion category and presenting stimuli of sufficient length to elicit the correct emotion categorization (Falagiarda & Collignon, 2019). As per results, besides the response at the general rate of presentation of all stimuli for both intact and scrambled sequences indicating common processes for all sounds, we also observed a response at the target frequency and its harmonics (Figure 2a,b). This could only occur if the brain was able to *discriminate* the target emotion from other emotion categories as well as *generalize* all sounds of the target emotion to one common category (Barbero et al., 2021). The significantly stronger responses to intelligible affective vocalizations when compared to scrambled sequences (Figure 2c,d), in conjunction with controlling of low-level features of the sounds indicate that the EEG emotion selective response is at least partially independent of low-level acoustic features and characterize a higher-order categorization process. Acknowledging the fact that the harmonicity in scrambled sounds was disrupted, the difference in EEG responses between intact and scrambled sequences was unlikely to arise due to differences in the harmonicity of our sounds since the frequency intact sounds had comparable HNR (Figure 1a).

We tested two types of sequences with either Fear or Happiness as the target emotion category repeated periodically at every third position (unbeknown to the participant). They expressed different topographies with part of their response spanning across few common right temporal channels suggesting the involvement of distinct yet overlapping neural substrates in emotion processing (Hamann, 2012; Johnstone et al., 2006; Mauchand & Zhang, 2022; Phan et al., 2002; Phillips et al., 1998). The significant response to the Fear category suggests the involvement of fronto-central and temporal areas, in line with neuroimaging studies indicating large scale networks involved in the processing of fearful expressions (Kober et al., 2008; Zhou et al., 2021). Further, the different scalp topographies found for Fear and Happiness suggest different generators for both processes and thus, can support the idea of different neural representations for categorization of different discrete emotions. In fact, previous research in vision that used that Fast Periodic Visual Stimulation (FPVS) also found responses to different facial emotions spanning different topographies (Dzhelyova et al., 2017; Leleu et al., 2018; Poncet et al., 2019). Although the arousal ratings of all discrete emotions were similar in our study, a possible reason for different topographies may be due to the difference in valence between Fear and Happiness (Figure 4d). However, valence does not seem to be a strong driver since we observed a strong oddball response when Fear is the target category and Fear was not deviant in terms of valence when compared to the other emotion categories presented in the sequence. Nevertheless, the mental states that support mechanisms of emotion processing can be abstract and highly dimensional and require further studies to disentangle which part of the processes linked to emotion discriminations are better conceptualized as categorical or dimensional (see Giordano et al., 2021; Hamann, 2012; Kragel & LaBar, 2016; Skerry & Saxe, 2015).

We also computed the time-course of the vocal categorization process. Both emotions elicited a significant response 300 ms post stimulus presentation, similar to studies that pointed out the late positive component (LPP) to emotional utterances (Frühholz et al., 2011; Jessen & Kotz, 2011). Additionally, the categorical response to Fear emerged in the brain as early as 135-175 ms post stimulus presentation. While previous literature has reported P200 time window to be modulated by emotional interjections and non-verbal vocalizations (Charest et al., 2009; Jessen & Kotz, 2011; Sauter & Eimer, 2010; Schirmer et al., 2013), it is debated whether these modulations are linked to the categorical nature of emotion expressions or to differences in arousal (Paulmann et al., 2013). Due to comparable affective dimensions (valence and arousal) of Fear with other categories in our stimuli sequence (Figure 4c,d), we speculate that the early ERP evoked by various vocalizations depicting Fear could be an early marker of its categorization, supporting recent evidence of the brain’s ability to represent discrete categories as early as <200ms (Giordano et al., 2021). In fact, many animal studies found primitive circuits for recognizing danger or fear preserved in mammals and in humans (LeDoux, 2012), making it possible to process fear faster than other emotions, notably due to potential contribution of subcortical pathways (Pessoa & Adolphs, 2010). Further, the absence of differences between intact and scrambled sequences before 100 ms supports the idea that the EEG frequency responses to both emotion categories are not elicited due to low-level acoustic properties of the sounds which are well known to modulate early ERP such as the N100 (Näätänen & Picton, 1987). Another interesting observation found in both frequency and time domain analysis (post 300 ms), was that the responses from four channels in the right temporal region (P6, TP8h, CP6 and T8h) were found to be significantly greater for intact sequences than scrambled for both Fear and Happiness conditions, putatively suggesting common processes for vocal emotion processing in these right temporal regions.

In summary, our research demonstrates categorical responses from the human brain to different discrete vocal emotions that are at least partially independent from processing of acoustic features. The frequency tagging method combined with EEG recordings has the advantage of obtaining such responses with a high signal-to-noise ratio and does not require the participant to overtly focus on the emotional content of sounds. Due to the high frequency resolution and temporal precision of EEG, we can expect the response at predefined frequencies that are adjustable with the experiment design, thus providing objectivity to the approach. Unlike many other ERP studies where emotion responses are compared to neutral causing confounds due to difference in stimuli acoustic features, this method provides a direct approach to obtain vocal emotion responses to discrete categories by including various heterogenous exemplars of each category. These characteristics make frequency tagging a valuable technique to study emotion categorization, suitable to use on populations that are more difficult to test (e.g. individuals with autism, infants etc) with classical paradigms such as ERP design studies and neuroimaging techniques.

## Acknowledgements

We would like to extend our gratitude to Anna Krzyżak for her help in EEG data acquisition.

## Notes

### Competing Interest Statement

The authors have declared no competing interest.

