## Supplementary Material for "Automatic brain categorization of discrete auditory emotion expressions"

Z-scores for all significant channels found as per the results section 3.1.2.

Table 1

| Channel | z-score | Channel | z-score | Channel | z-score |
| --- | --- | --- | --- | --- | --- |
| Fear Intact > Scrambled FDR corrected | | | | | |
| Cz | 2.4133 | B30 | 2.6852 | D1 | 3.5852 |
| I1 | 2.2329 | FC4h | 2.7230 | FCC1h | 5.1274 |
| Oz | 2.8293 | FCC2h | 2.3914 | FFC3h | 5.0419 |
| I2 | 2.5596 | FCC2 | 4.5105 | FFC3 | 5.4294 |
| POI2 | 2.3645 | FFC2 | 3.7994 | FT7 | 2.7384 |
| P10 | 3.3620 | F2 | 3.0374 | FT7h | 3.1347 |
| P6 | 2.2327 | AFF4h | 3.1518 | FC5 | 2.7468 |
| TP8 | 4.2504 | AFFz | 3.4333 | FC3h | 4.4714 |
| TP8h | 5.1871 | Fz | 2.9558 | C1 | 3.3253 |
| CP6 | 3.1889 | FFCz | 3.3953 | C5 | 2.8902 |
| T8h | 4.4285 | FCz | 4.7923 | T7h | 2.4928 |
| T8 | 3.4197 | FFC1 | 4.7615 | T7 | 3.1867 |
| FT8 | 4.8867 | F1 | 4.3491 | TP7 | 3.9254 |
| FT8h | 5.3219 | AFF1 | 3.5015 | TP7h | 4.7200 |
| Happy Intact > Scrambled FDR corrected | | | | | |
| CPP3 | 3.0811 | CP6h | 2.7744 | AFF5 | 4.3399 |
| C2h | 2.9064 | C4 | 3.3134 | F5 | 2.6541 |
| P6 | 2.7621 | C6h | 4.8022 | F7 | 3.0511 |
| TP8 | 3.3129 | C6 | 2.9586 | P7 | 3.5407 |
| TP8h | 2.9090 | T8h | 3.6893 |  |  |
| CP6 | 3.6049 | AF7 | 3.0840 |  |  |
| Fear Intact > Happy Intact FDR corrected | | | | | |
| Cz | 3.6662 | FCC2 | 4.9567 | FC5 | 2.8393 |
| CCPz | 3.3739 | FFC2 | 6.5864 | FC3 | 4.4699 |
| CPz | 3.4205 | F2 | 3.5489 | FC3h | 3.2799 |
| I1 | 2.4174 | Fz | 2.5007 | C1 | 3.8339 |
| I2 | 3.5345 | FFCz | 5.6050 | C1h | 2.9282 |
| POI2 | 2.9950 | FCz | 6.5102 | CCP1h | 6.3831 |
| O2 | 2.7675 | FFC1 | 6.4061 | C3h | 2.9730 |
| P2 | 2.2944 | F1 | 5.0717 | C3 | 2.2431 |
| C2h | 3.1041 | FCC1h | 4.1906 | C5 | 2.8818 |
| C4 | 2.4353 | FCC1 | 4.8892 | T7 | 3.4979 |
| FT8 | 2.8416 | FFC3h | 3.2197 | TP7 | 3.0771 |
| FT8h | 4.7199 | FFC3 | 3.0675 | P9 | 3.6068 |
| FCC2h | 3.9286 | FT7 | 2.5508 |  |  |
